# Assessing genome conservation on pangenome graphs with PanSel

**DOI:** 10.1101/2024.04.26.591236

**Authors:** Matthias Zytnicki

## Abstract

1

**Motivation:** With more and more telomere-to-telomere genomes assembled, pangenomes make it possible to capture the genomic diversity of a species. Because they introduce less biases, pangenomes, represented as graphs, tend to supplant the usual linear representation of a reference genome, augmented with variations. However, this major change requires new tools adapted to this data structure. Among the numerous questions that can be addressed to a pangenome graph is the search for conserved or divergent genes.

**Results:** In this article, we present a new tool, named PanSel, which computes a conservation score for each segment of the genome, and finds genomic regions that are significantly conserved, or divergent.

**Availability:** PanSel, written in C++11 with no dependency, is available at https://github.com/mzytnicki/pansel.

## 2 Introduction

Genome diversity has long been studied for genetic diseases, genome improvement of species of agricultural interest, among others. With the advent of new, long, almost errorless, reads, it is now possible to produce several telomere-to-telomere haplotypes for complex eukaryotes. These haplotypes, if they include the diversity of the population of interest (usually, a species), may be compared, leading to pangenomic studies [1]. A pangenome can be efficiently stored into a dedicated data structure, which is usually called a variation graph [6]. This graph contains so-called *segments*, which are sequence chunks found in at least one haplotype, and *paths*, which are lists of segments (possibly in reverse-complement) representing haplotypes. A given segment, if it belongs to several paths, represents a homologous sequence, shared by the haplotypes.

Although pangenomic studies have been published for almost two decades in prokaryotes, they are relatively new for multi-cellular eukaryotes, and changing the framework from a genome supplemented with variations to a variation graph is no simple task. Notably, it requires adapting all the tools which were developed for the previous framework, and there are numerous. It is for instance crucial to be able to assess the conservation, or the divergence, of genome *loci*, *i.e.* sequences that evolve slower (or faster) than the expected natural drift. Several methods have been developed for inter-species analyses with a reference genome for each organism, and they include phastCons [17], GERP and GERP++ [2], and phyloP [14]. They use a multiple sequence alignment, and a phylogeny, in order to estimate the length of the branches of the tree at each position of the genome alignment. These metrics are then widely used, available in Genome Browsers [15], and can be used to check whether a gene of interest is conserved through evolution.

To date, there is no such method for intra-species conservation studies, and inter-species tools cannot be be used as is. For instance, the phylogenetic tree is usually not known, or even not defined, between individuals of the same species, because inbreeding may be frequent. Some methods do find conserved regions, usually using *k*-mers, such as Corer [16], or [12], and cluster them into “cores” or “pan-conserved segment tags”. Yet, finding divergent regions, or assessing the conservation of a gene of interest, is not possible.

Our aim is to provide a simple tool, that could be applied by consortia that produce variation graphs, in order to assess the conservation of each point of the genome, so that users could check whether their genes of interest are under selection pressure, or accumulating variations.

We wanted to leverage the variation graph, because it is now the standard model for representing diversity. Yet, the base-level multiple alignment —which tentatively puts in the same column the nucleotides that evolved from the common, ancestor nucleotide— is not available in the variation graph. Finding a conservation value for each nucleotide is thus not directly applicable. So, we depart from the base-level resolution, which is not easily accessible using a pangenome graph. Instead, we use a sliding windows of fixed size (*e.g.* of size 1kb), which is enough to detect conserved and divergent genes. Second, we estimate the diversity by computing the Jaccard index (as defined by ODGI [8]) between each pair of haplotypes in each window. Last, significantly under- and over-conserved windows are detected by fitting a mixture model.

## 3 Implementation

PanSel is a C++ tool that reads a GFA file representing one chromosome (it thus cannot, in its current version, deal with chromosomal translocations). Given a window size *s*, it tries to detect conserved segments (named *boundary segments* hereafter) distant by *s* nucleotides on the reference path. For each pair of boundary segments, it extracts the sub-path for each haplotype. The (weighted) Jaccard index [8] is then computed.

Since the Jaccard index belongs to the [0, 1] interval, we transform it R_+_ using the *−* log function. The distribution is then fitted with a simple mixture model. The most conserved part follows a Gaussian distribution. However, we observe a heavy tail on the right-hand side of the distribution, which represents divergent sequences, and is not compatible with a Gaussian distribution. We model it using a log-normal distribution, and finally fitted the whole distribution with the average of the two previously found distributions. A more detailed description of the implementation, and the fit, is given in Supplementary Section 1.

## 4 Results

We applied PanSel with three sliding window sizes (1000, 10,000, and 100,000) on the Draft Human Pangenome [13]. We took the 2022-03-11 release, computed by MiniGraph-Cactus [10]. We tested another tool, PGGB [3]. We noted that its graphs are more fragmented, *i.e.* chromosomes are split, most likely in the smoothxg step. As a result, fewer paths connect a boundary segment to the next segment, and results are less reliable, especially in complex regions. We thus discarded PGGB, although the GFA produced by this tool is correctly parsed. In our computer (Intel® Xeon® CPU E5-2687W v4 @ 3.00GHz running Debian x86 64 6.1.76-1), PanSel took less than 25 minutes, and not more than 20GB RAM, for each chromosome (run on a single thread per chromosome).

### 4.1 Comparison with other annotations

First, we discarded the regions that overlapped with known gaps (see Supplementary Section 1.6). We then divided the regions into 8 categories, with the most conserved regions being in the first bin, and the most divergent regions being in the last bin. For the sake of space, we will only present results for a window size of 1kb, but results for other windows sizes follow the same trend, and are presented in Supplementary Section 3.1.

We compared each bin with several available annotations. First, we counted the number of structural variants that overlapped with each bin. As seen in Figure 1 (first row), the number of variants increases with the bin number, and the final bin overlaps with significantly more known variants.

**Figure 1:**
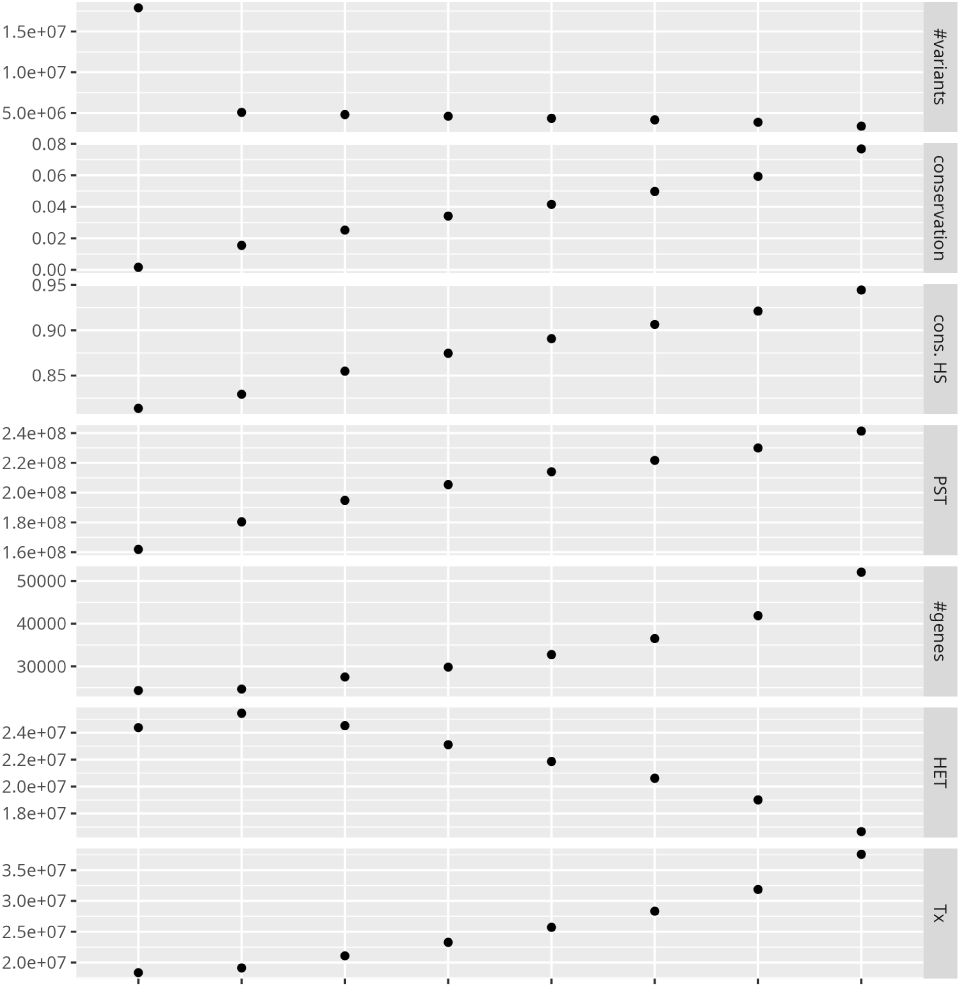
Each column represents a bin. The bins on the left include the most conserved regions, whereas the ones on the right include the most divergent regions. For each bin, we computed the number of overlapping structural variants (first row), the average vertebrate conservation, the average human conservation, the number of pan-conserved segment tags, the exonic coverage, the predicted heterochromatin coverage, and the predicted actively transcribed coverage (last row).

We computed the average conservation score, provided by PhyloP [14] using 100 vertebrates, on each bin. The second row of Figure 1 shows that the conservation decreases with the bin number. This confirms that genomic regions that are more conserved in human are also more conserved in vertebrates.

We also computed the PhastCons [17] score (which took several days to complete), based on a human pangenome multiple alignment, and provided the average score in the third row. The number of pan-conserved segment tags [12] is given in the fourth row. The different methods provide concordant results. We tried to use Corer [16], but it did not complete in four days with 80 CPUs. We compared the bins with the location of known protein coding exons, provided by GenCode [5], and computed the number of exonic nucleotides found in each bin. As seen in the fifth row of Figure 1, conserved regions contain more coding exons, as expected.

Last, we extracted the annotation computed by ChromHMM [4]. Briefly, ChromHMM splits the genome into 100 categories [18], based on ChIP-Seq data. These categories are associated with different states of the chromatin (*e.g.* “actively transcribed”, or “Tx”). Each state may have different levels (*e.g.* there are eight “Tx” levels), and we merged these levels in this study. Here, we focused on the “Tx” and the “HET” (for heterochromatin) states, but all the results can be found in Supplementary Section 3.2. As seen in the sixth row of Figure 1, heterochromatin-rich regions are more divergent. The last row shows that actively transcribed regions are more conserved, as expected.

### 4.2 Study of a particular gene

We wanted to showcase the use PanSel on the human genes. We extracted the coding genes that co-localized with the most divergent regions. Among them, we found *ANKRD30A*, *BRF1*, *NBPF20*. The example of the latter gene is presented in Figure 2, while the others are presented in Supplementary Data, Section 3.3.

**Figure 2:**
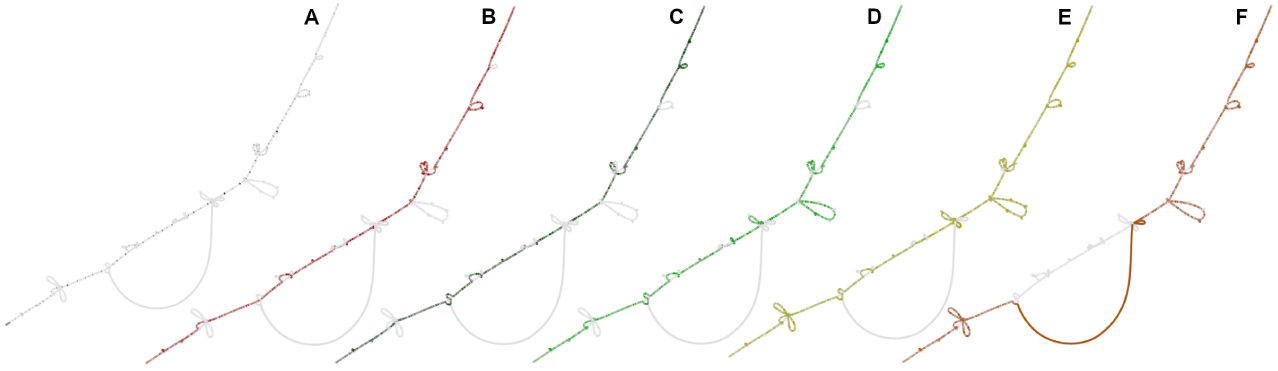
Bandage [19] captures of different haplotypes of *NBPF20*. A: Grey lines show all the haplotypes, and black dots show exons of *NBPF20*, as annotated in GRCh38. B: The highlighted path represents the reference assembly GRCh38. C: The highlighted path represents the telomere-to-telomere assembly CHM13. D-F: The highlighted paths represent haplotypes of NA21309.1 (Maasai), HG00735.2 (Puerto Rico), and HG02559.2 (Barbados), respectively.

*NBPF20* belongs to the neuroblastoma breakpoint family [9, 7], which recently expanded, especially in humans. Some genes may thus undergo neo- or sub-functionalization, and accumulate variations. Figure 2 shows the human pangenome graph, restricted to this gene. Figure 2A, we highlighted in black the exons of the gene (as annotated in GRCh38). Figure 2B and C, we highlighted the GRCh38 and CHM13 paths respectively. The last three subfigures show haplotypes of different people. These figures pinpoint the wealth of SVs in the diversity. Some of them may alter the transcript: some exons are skipped and are thus missing in the final transcripts; some exons may also be added, since new genetic material is present. SVs in this gene family, which include higher order repeats, are well documented, and may be linked with cognitive disabilities, among others. Proving that this gene is evolving rapidly in humans, taking into account both SNPs and SVs, has not been done before, and highlights the usefulness of PanSel.

### 4.3 Conserved and divergent regions

Another use of PanSel is the study of regions that are significantly conserved and divergent. We used a p-value of 5% for a window size of 10,000 (a window size of 1,000 is usually too small to capture the full extent of a gene). We found about 79Mbp and 162Mbp of conserved and divergent regions, respectively. Figures 9 to 11 in Supplementary Data show the distribution of conserved and divergent regions on the autosomes. As expected, divergent regions accumulate near the telomeres, while the convergent regions are located inside chromosomes.

We then extracted the overlapping genes (4,861 and 12,286 genes in conserved and divergent regions, respectively), and looked for enriched gene ontologies using gProfiler [11].

Full results are provided in Supplementary Section 3.3. Briefly, conserved regions are related to very general molecular functions (*e.g.* protein binding), biological processes (*e.g.* multicellular organism development) or cellular compartment (*e.g.* nucleoplasm).

Divergent regions are related to more specific molecular functions (*e.g.* antigen binding), biological processes (*e.g.* adaptive immune response) or cellular compartment (*e.g.* immunoglobulin complex), which are related to immunity.

This would suggest that basic molecular functions are located in conserved regions, whereas immunity related genes are located in divergent regions, as expected.

## 5 Acknowledgements

We are grateful to the genotoul bioinformatics platform Toulouse Occitanie (Bioinfo Genotoul, https://doi.org/10.15454/1.5572369328961167E12) for providing computing and storage resources. We also deeply thank the Reviewers for very insightful suggestions.

## 6 Supplementary data

Supplementary data are available at *Bioinformatics Advances* online.

## 8 Data availability

The pipe-line used to produce the results can be retrieved from https://github.com/mzytnicki/pansel_paper.

## A Description of the method

### A.1 GFA parsing

PanSel reads the S, P, and W lines of the GFA file. All the other lines are skipped. It stores the lengths of the segments (but not the nucleotidic sequences), and the paths (as lists of segments). In some GFA files, variations graphs have been enriched with additional variations, some of them being named_MINIGRAPH_.sXXXX (where X is a number) by the MiniGraph-Cactus pipe-line. These lines are skipped.

### A.2 Conserved segments

In this step, PanSel finds all the segments that are included in all paths. These segments represent conserved sequences. The number of haplotypes can be provided by the user. Alternatively, PanSel can detect it. The *conserved* segments that are included in all the paths are then stored.

### A.3 Sliding windows

PanSel then follows the path of the reference haplotype (provided by the user). It scans each segment, which can be represented as a genomic interval [*a, b*] on the reference haplotype. It stops at the first conserved segment, *A* representing interval [*a, b*]. This is the first boundary segment. From there, it tries to find the next boundary segment, *B*, such that the distance between *A* and *B* is approximately *w*, the size of the user-defined sliding window. More formally, we are looking for, *B*, representing interval [*c, d*], such that *c − b ≤ w ≤ d − a*.

There may be no such boundary segment: the position of the ending sliding window may contain variations. In this case, we look for the next boundary segment, and record that the size of the sliding window will be larger than expected.

We then proceed to the next interval.

### A.4 Computing the weighted Jaccard index

We extract the sub-paths of each haplotype between each pair of selected, consecutive boundary segment. Given two sub-paths, stored as two sets of segments *A* and *B*, the Jaccard index is the ratio between the size of the intersection of the segments, divided by the size of union:

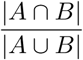

However, some segments are longer than others, and a SNP should “weight” less than a structural variation of, say, 100 base pairs. Following the method defined in ODGI, we use the following formula as the weighted Jaccard index (where *s* is a segment, and *|s|* is its length):

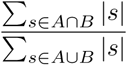

For each sliding window, we compute the average weighted Jaccard index between all pairs of sub-paths.

### A.5 Fitting the average weighted Jaccard index

We first compute the distribution of the average weighted Jaccard index per region. Since the index is in the [0, 1] interval, we translate it to the R_+_ using the *−* log transformation. We compute the mode *m*, and extract the region corresponding to *x ∈* [0*, m*]. We add the symmetric of this distribution, and fit a normal distribution. We also fit the whole distribution with a log-normal, and plot the average of the two distributions.

The *p*-value can then be computed directly from the mixture model. In order to find the significantly conserved (resp. divergent) regions, we take the extreme 5% of the normal (resp. log-normal) distribution. The threshold can be modified by the user.

The fit are plotted in Figures 3, 4, and 5.

**Figure 3:**
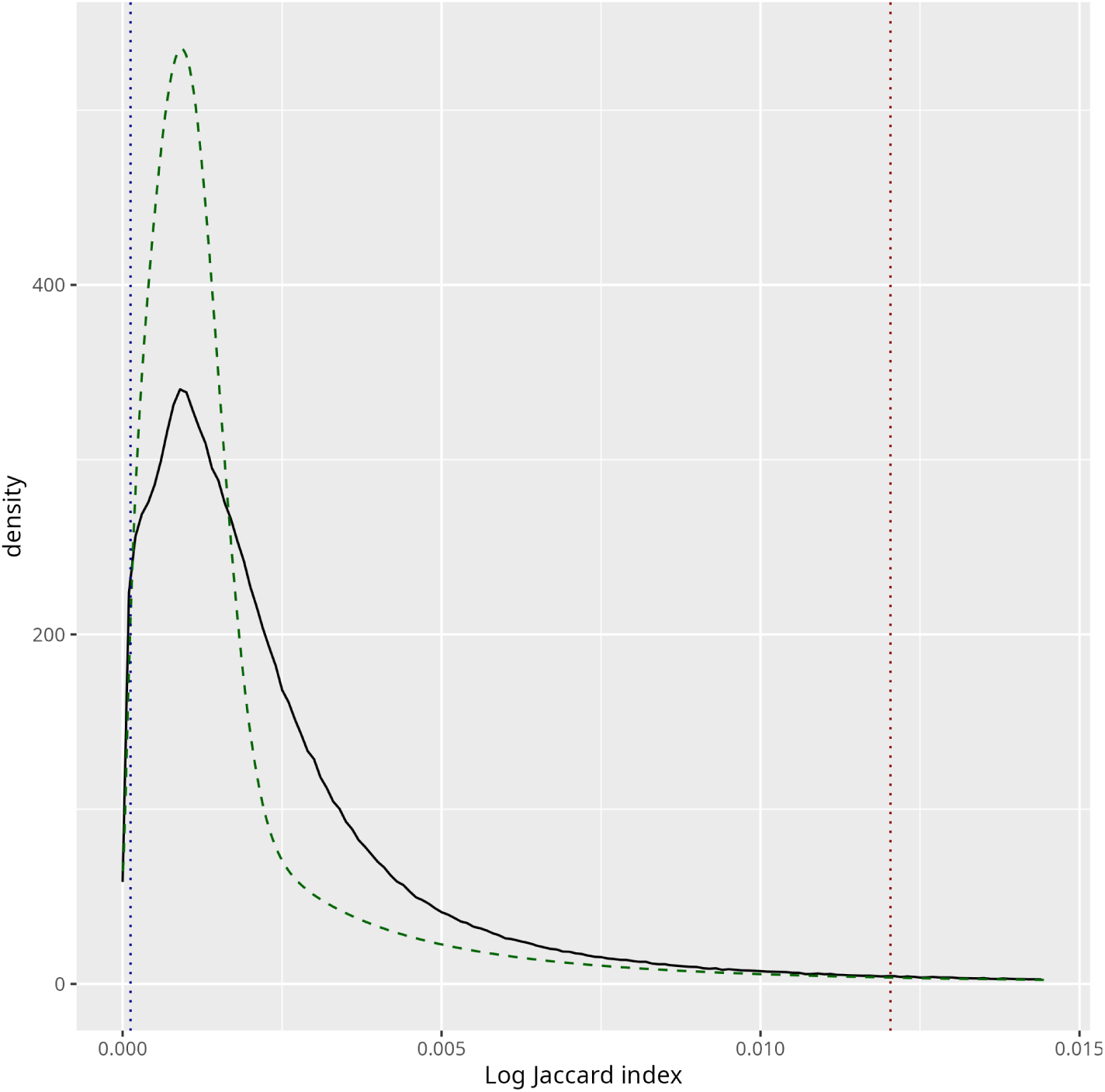
Distribution of the average weighted Jaccard index, in solid black line, for a window size of 1000. The fit is in dashed green, the threshold for significantly conserved (resp. divergent) regions is in dotted blue (resp. red).

**Figure 4:**
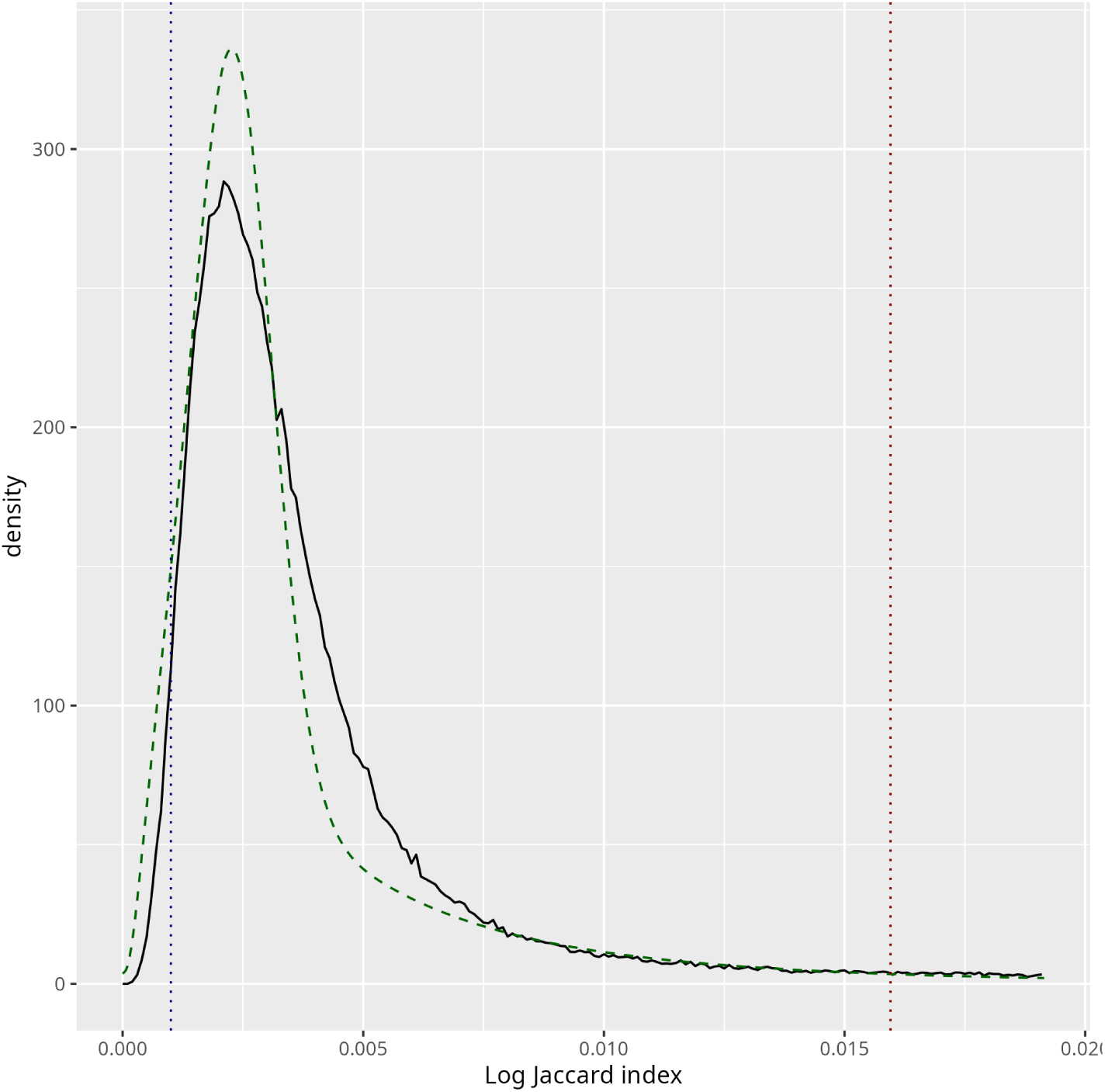
Distribution of the average weighted Jaccard index, in solid black line, for a window size of 10,000.

### A.6 Splitting data to different conservation strata

The script divideRegions.py (see the corresponding Github page) reads the output of Pansel. It first removes regions that are too large (1.5 times the size of the bin): these correspond to regions where it is not possible to find anchors. Then, it also discards regions that co-localize with gaps (as given by the annotation consortium). Then, it ranks the remaining regions based on the Jaccard index, and splits them into strata of (roughly) identical sizes.

**Figure 5:**
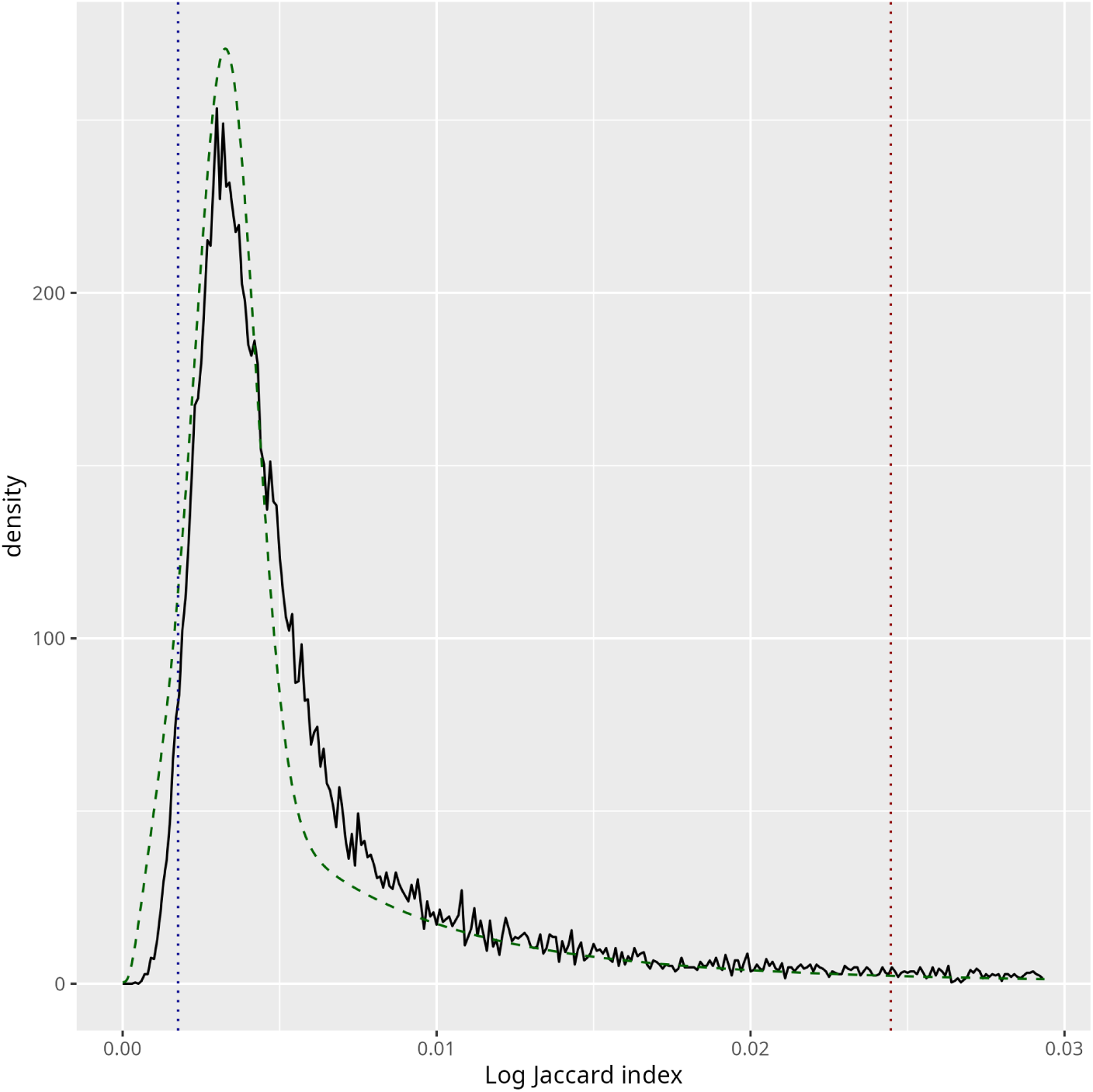
Distribution of the average weighted Jaccard index, in solid black line, for a window size of 100,000.

## B Data used

### B.1 Pangenome graphs

The pangenome graphs were downloaded from the Human Pangenome initiative: https://s3-us-west-2.amazonaws.com/human-pangenomics/index.html?prefix=pangenomes/scratch/

We chose the 2022 03 11 minigraph cactus release, and downloaded one vg file per chromosome.

Each file was then converted to GFA format using vg view.

We chose GRCh38 as a reference, since many annotation are available.

### B.2 Genome annotations

- Assembly gaps: https://hgdownload.soe.ucsc.edu/goldenPath/hg38/bigZips/latest/ hg38.agp.gz
- List of structural variants from dbVar: https://ftp.ncbi.nlm.nih.gov/pub/dbVar/data/Homo_sapiens/by_assembly/ GRCh38/vcf/GRCh38.variant_call.all.vcf.gz
- 100 vertebrate conservation using PhyloP: https://hgdownload.soe.ucsc.edu/goldenPath/hg38/phyloP100way/ hg38.phyloP100way.bw
- Genome annotation: https://ftp.ebi.ac.uk/pub/databases/gencode/Gencode_human/release_ 44/gencode.v44.annotation.gtf.gz
- ChromHMM annotation: https://public.hoffman2.idre.ucla.edu/ernst/2K9RS/full_stack/ full_stack_annotation_public_release/hg38/hg38_genome_100_segments. bed.gz
- Multiple alignent of human genome in HAL format (used by PhastCons): https://s3-us-west-2.amazonaws.com/human-pangenomics/pangenomes/ scratch/2022_03_11_minigraph_cactus/hprc-v1.1-mc-grch38-full. hal
- Pan-conserved segment tags: https://dna-discovery.stanford.edu/publicmaterial/datasets/pangenome/ pst-31mer.grch38.bed.gz

## C Results

### C.1 Scores for the other window sizes

This part gives the same results as Figure 1 in the main document, with different window sizes.

#### C.1.1 Window size of 10k

For 10k window size, results are very similar to 1k size.

**Figure 6:**
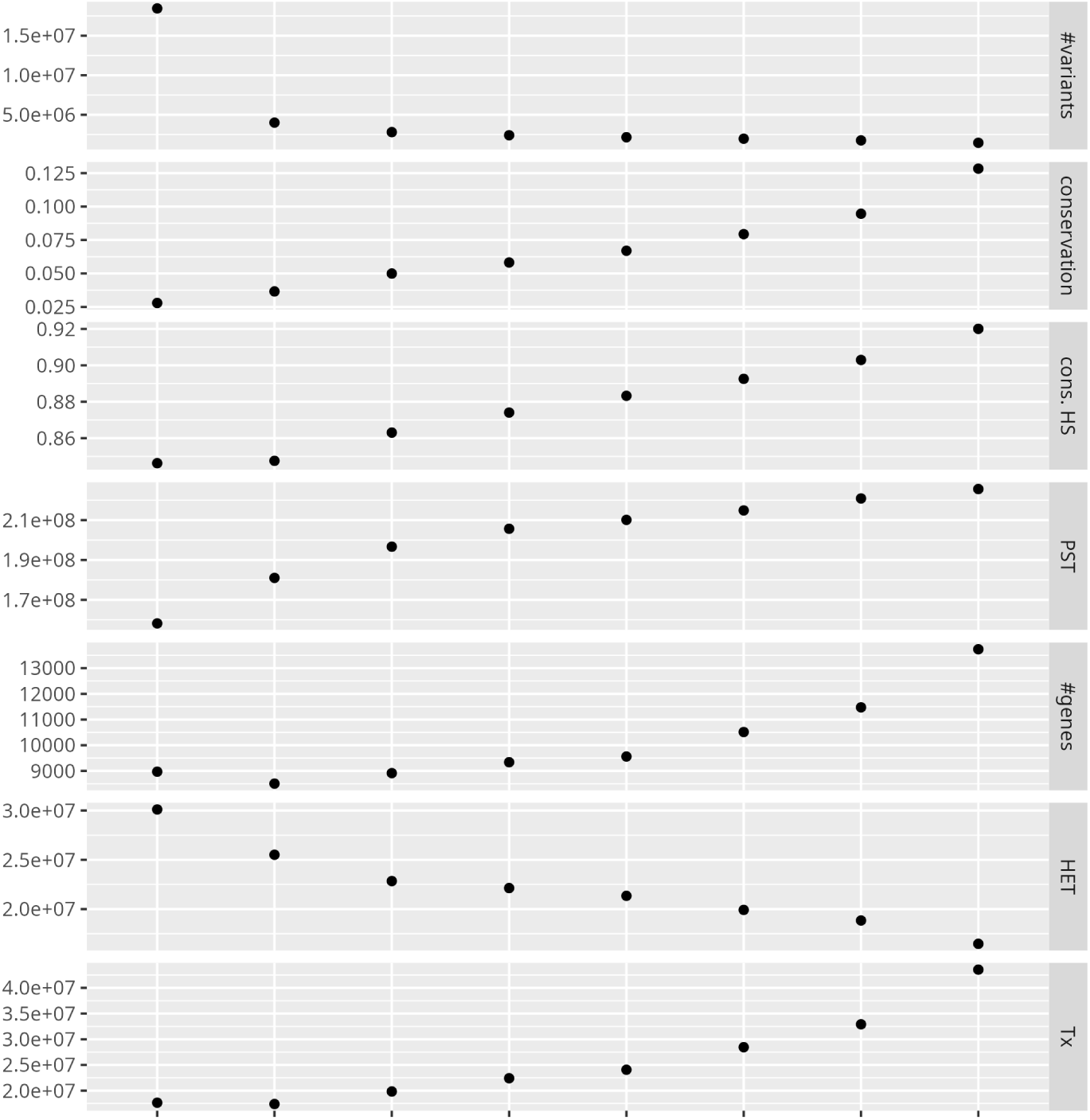
Results for window size of 10k. See Figure 1 in the main document for a detailed legend.

#### c.1.2 Window size of 100k

At 100k, the signal is less clear, probably because windows are too large, and include very diverse types of regions.

**Figure 7:**
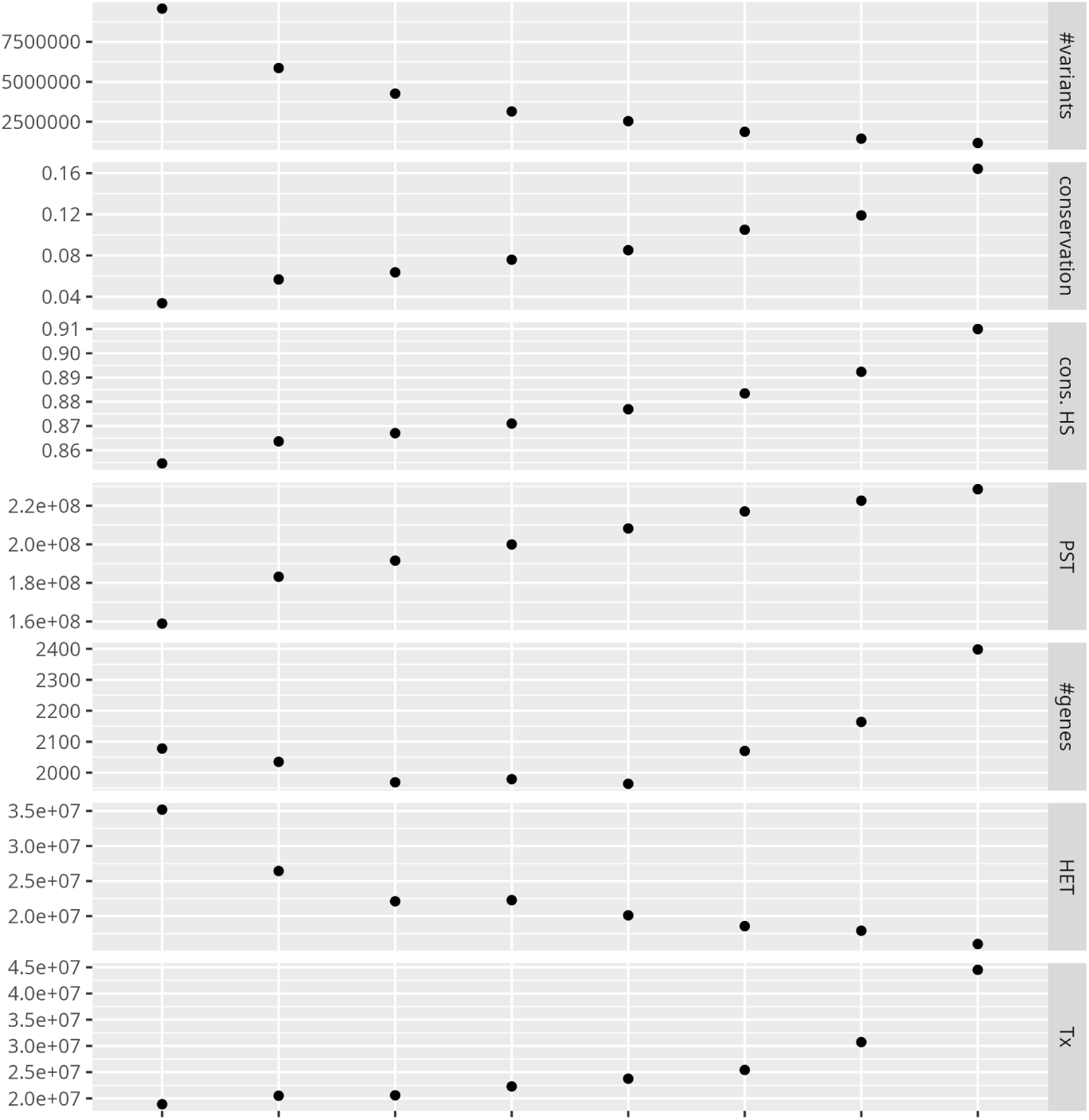
Results for window size of 100k. See Figure 1 in the main document for a detailed legend.

### C.2 ChromHMM full results

This part gives the coverage for all ChromHMM categories. The categories are:

- Acet: acetylations
- BivProm: bivalent promoters
- DNAse: DNase I hypersensitivity
- EnhA: active enhancers
- EnhWk: weak enhancers
- GapArt: assembly gaps and alignment artifacts
- HET: heterochromatin
- PromF: Flanking promoters
- Quies: repressive or inactive
- ReprPC: polycomb repressed
- TSS: transcription start site
- Tx: strong transcription
- TxEnh: transcribed enhancers
- TxEx: transcript exons
- TxWk: weak transcription
- ZNF: zinc finger

#### C.2.1 Window size of 1k

Figure 8 clearly shows that repressed states, including HET, Quies, and ReprPC, tend to co-localize with divergent regions. It is also the case for Acet states, which is a weak mark of promoter or enhancers.

**Figure 8:**
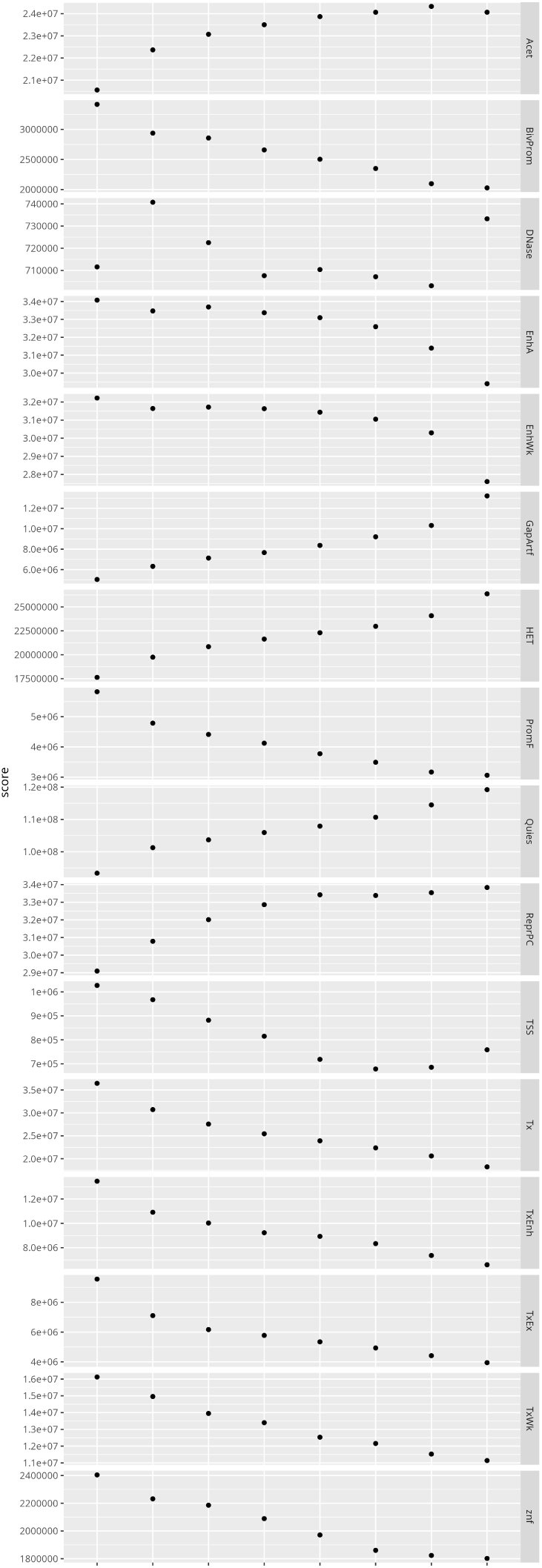
Results for window size of 1k.

On the other hand, active states, including BivProm, EnhWk, PromF, TSS, TxEnh, TxEx, TxWe, and ZNF, tend to co-localize with conserved regions.

The DNase state does not seem to follow any clear trend.

#### C.2.2 Window size of 10k

Results for a window size 10k are similar to the previous ones. The correlation, however, seems slightly less clear, probably because larger windows include different types of regions.

**Figure 9:**
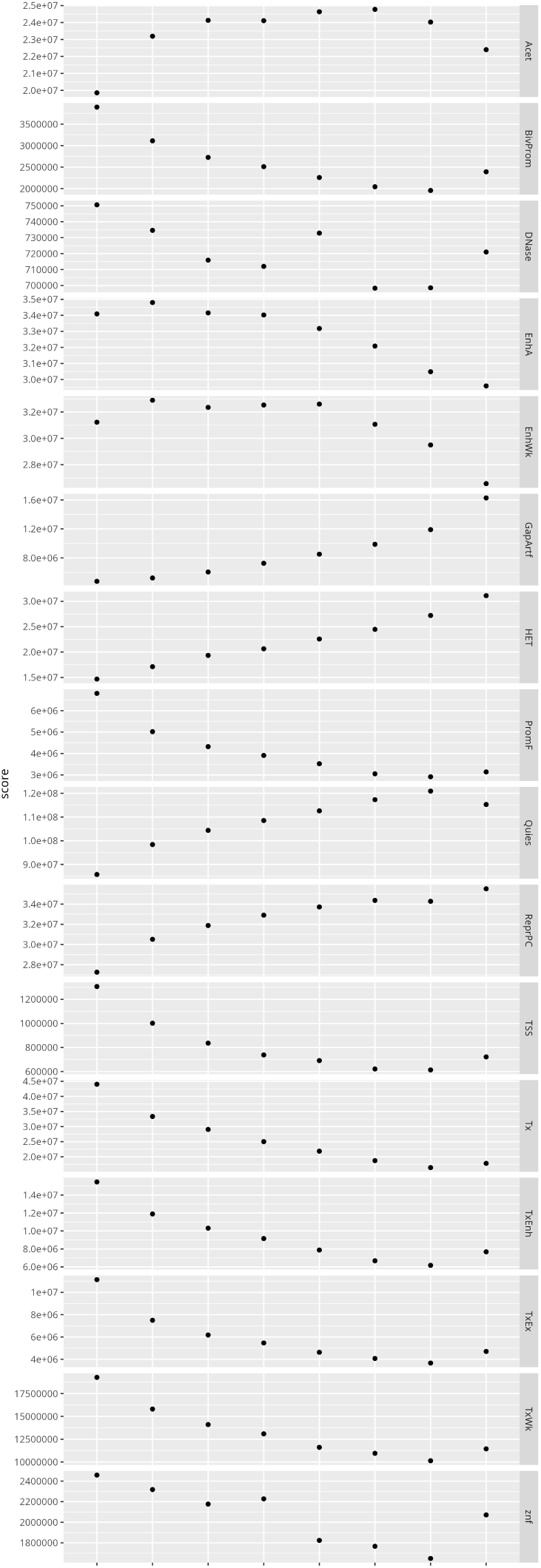
Results for window size of 10k.

#### C.2.3 Window size of 100k

Results for a window size 100k are similar to the previous ones, but even less clear.

**Figure 10:**
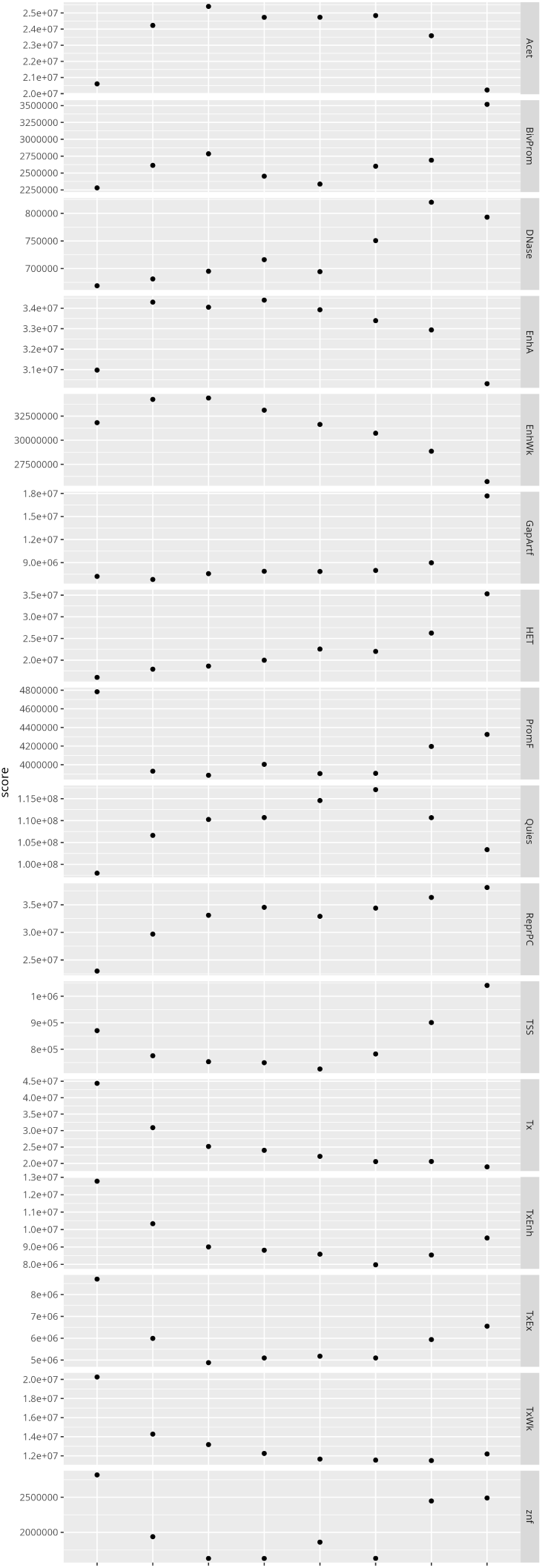
Results for window size of 100k.

### C.3 Study of particular genes

#### C.3.1 ANKRD30A

Figure 11 shows the gene *ANKRD30A* (Ankyrin Repeat Domain 30A), which has been associated with breast cancer.

**Figure 11:**
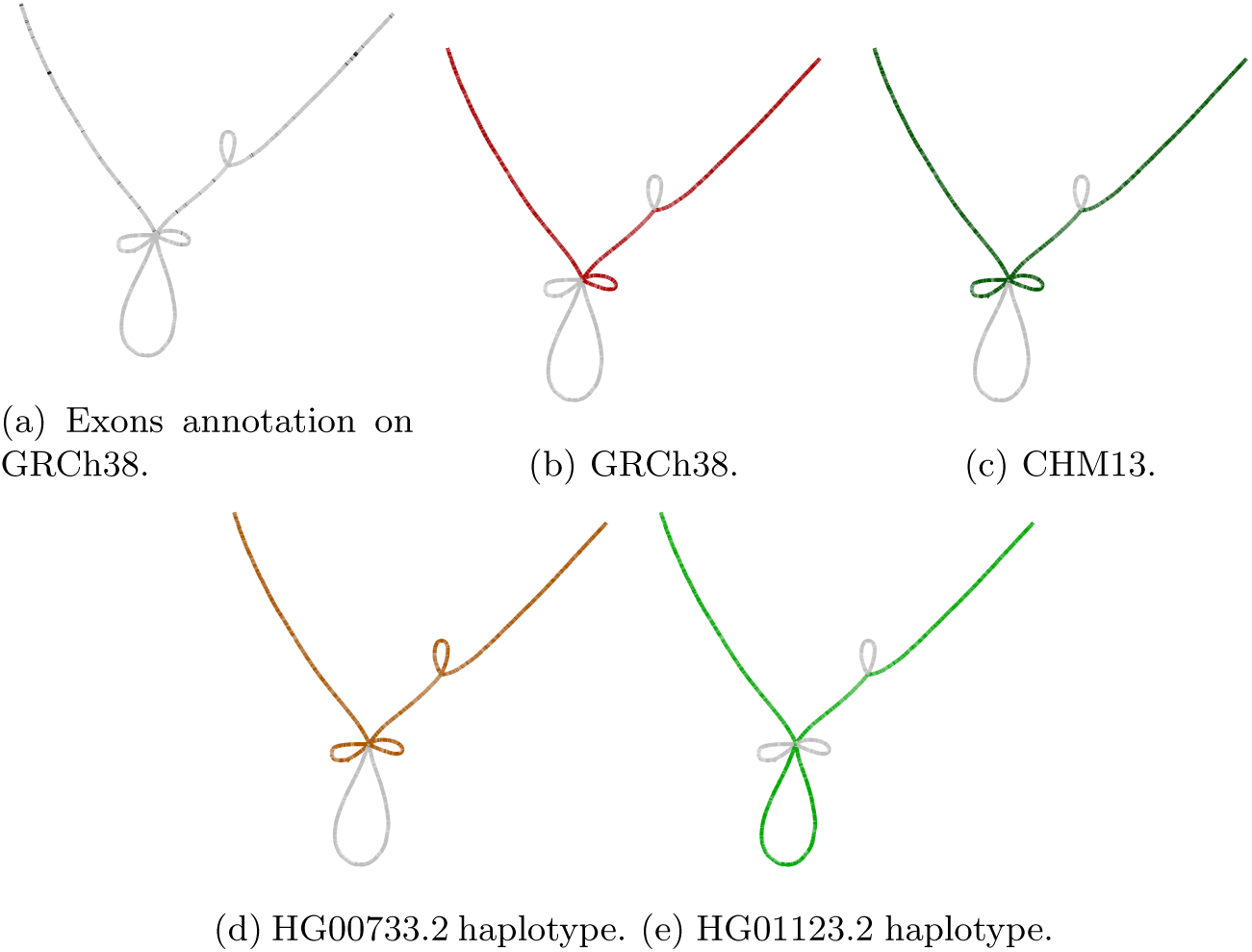
*ANKRD30A*: exon annotation on GRCh38, and different assemblies and haplotypes.

##### C.3.2 BRF1

Figure 12 shows the gene *BRF1*, which is one of the subunits of the RNA polymerase III transcription factor complex.

**Figure 12:**
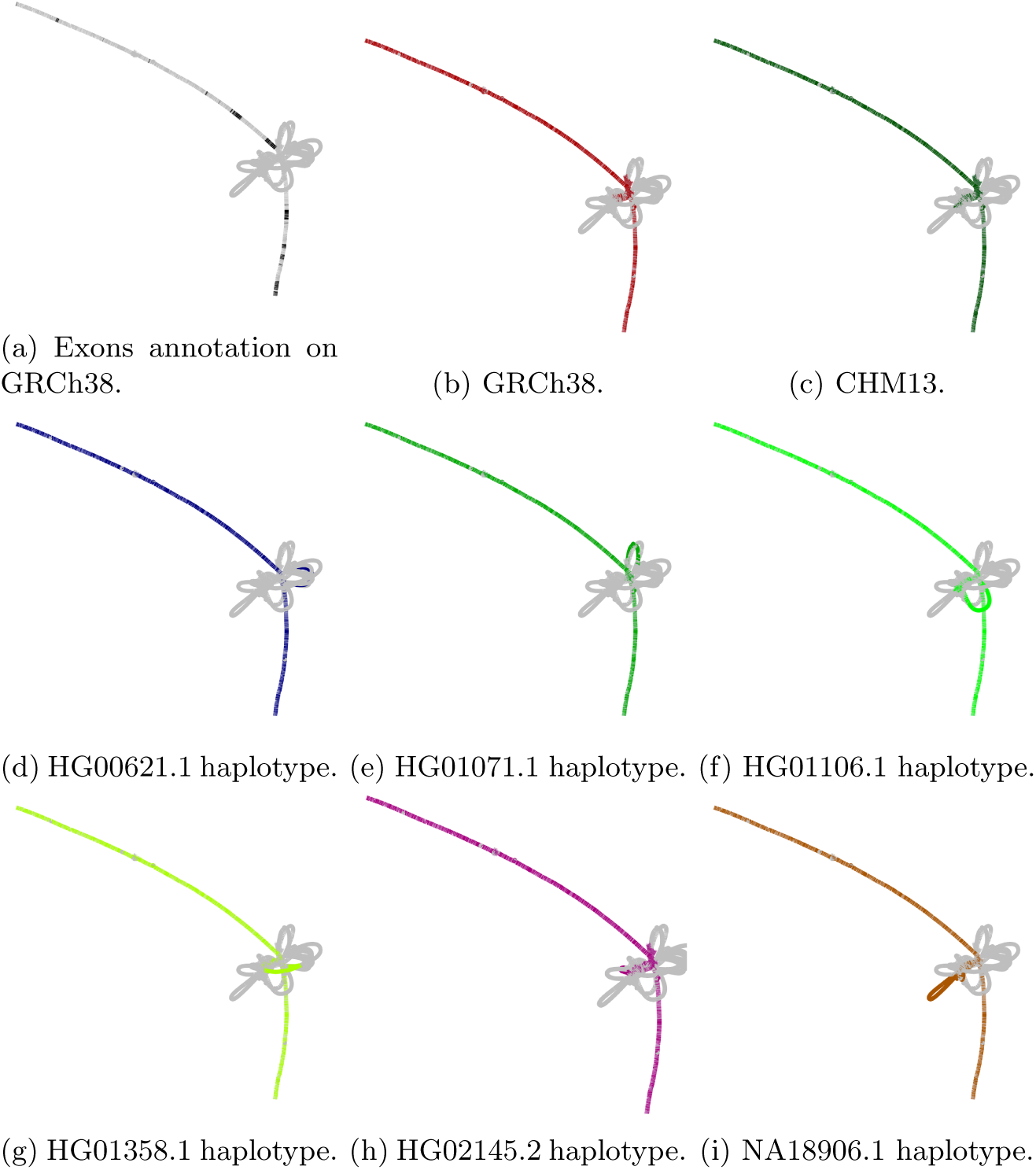
*BRF1* : exon annotation on GRCh38, and different assemblies and haplotypes.

### C.4 Density of conserved and divergent regions

This section provides the number of divergent and conserved regions along the chromosomes. The numbers on the *y*-axis gives the number of significant regions within 1Mbp. Results are given for the different bin sizes.

#### C.4.1 Window size of 1k

**Figure 13:**
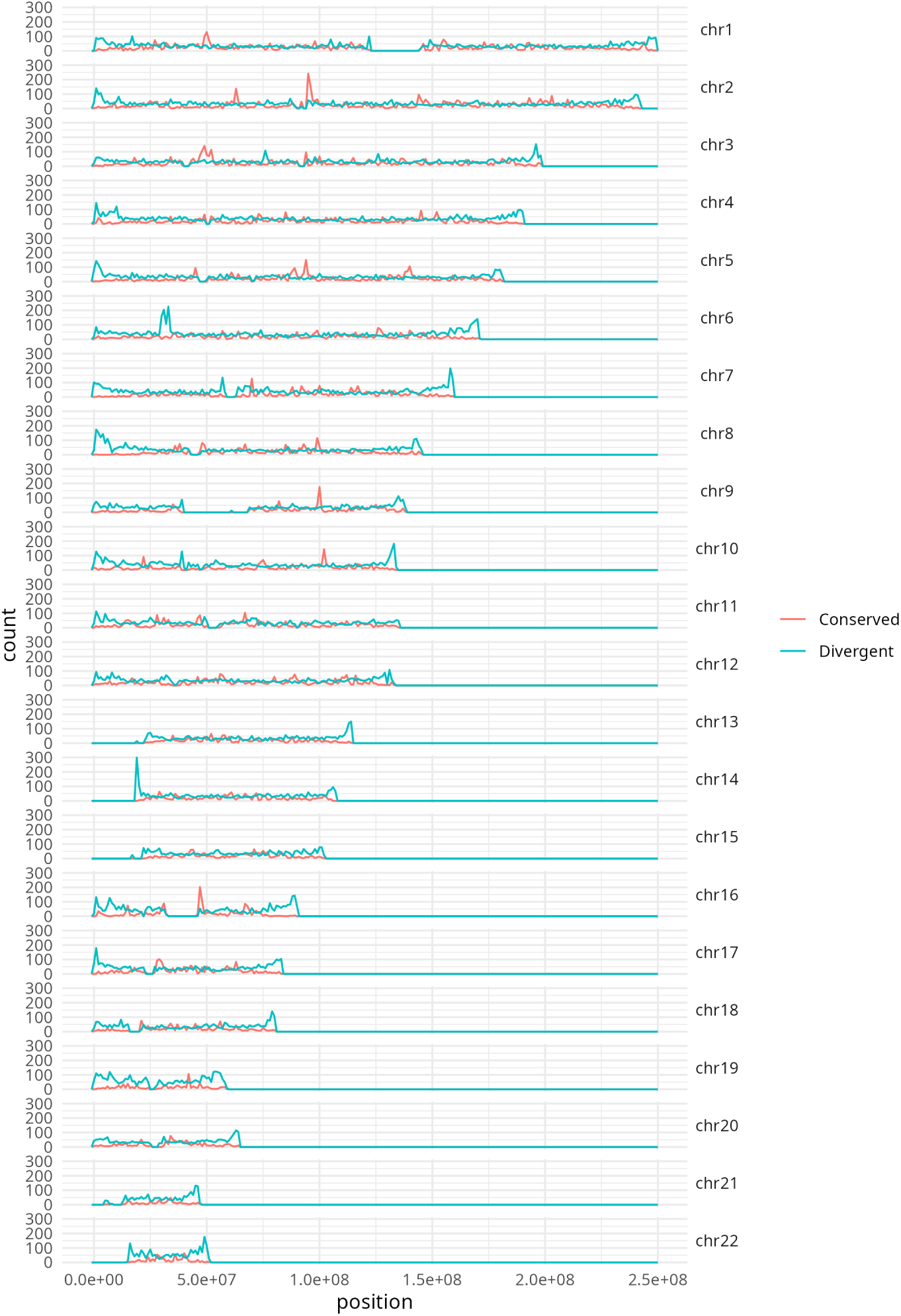
Density of conserved and divergent regions for bin size of 1k.

#### C.4.2 Window size of 10k

**Figure 14:**
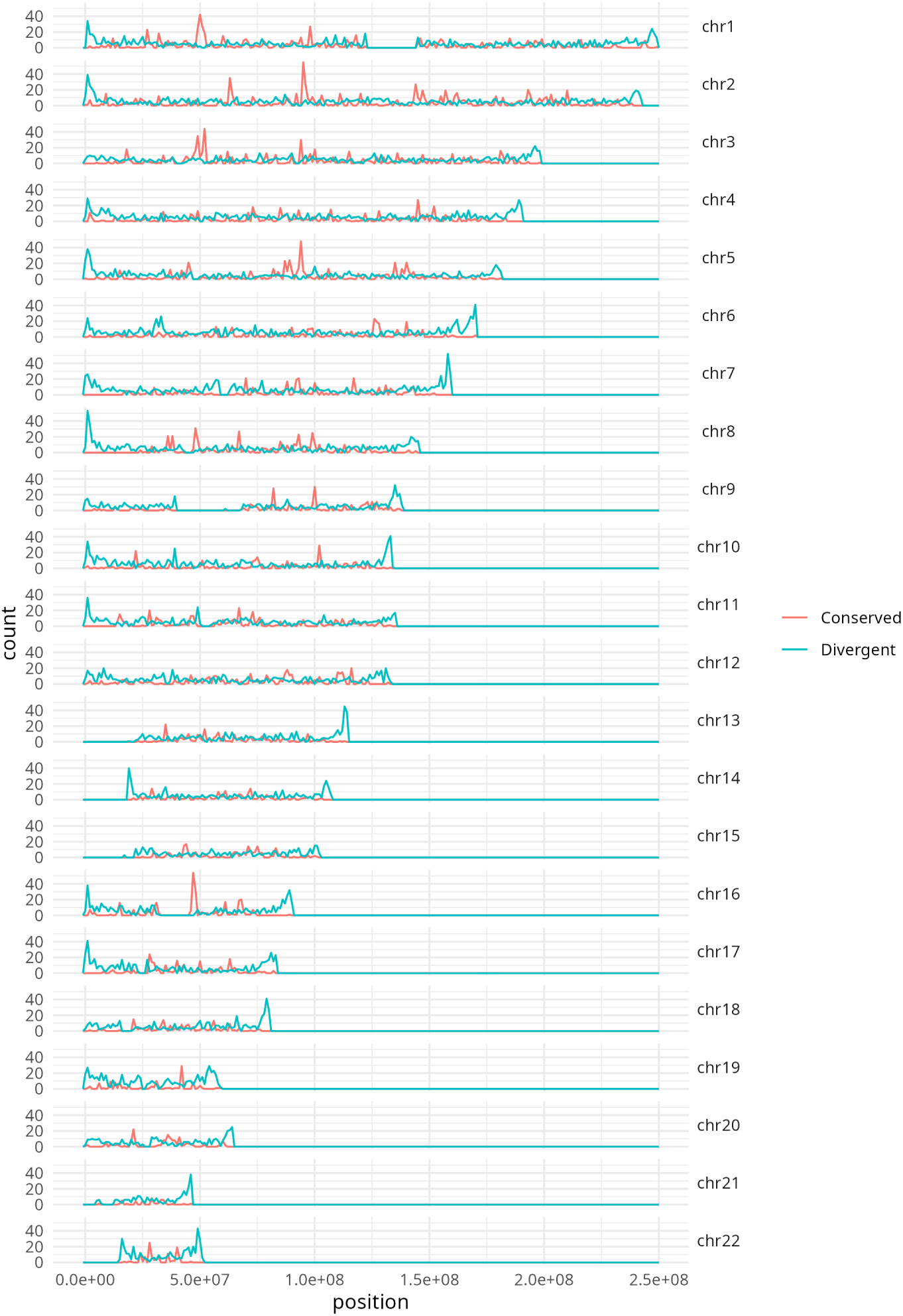
Density of conserved and divergent regions for bin size of 10k.

#### C.4.3 Window size of 100k

**Figure 15:**
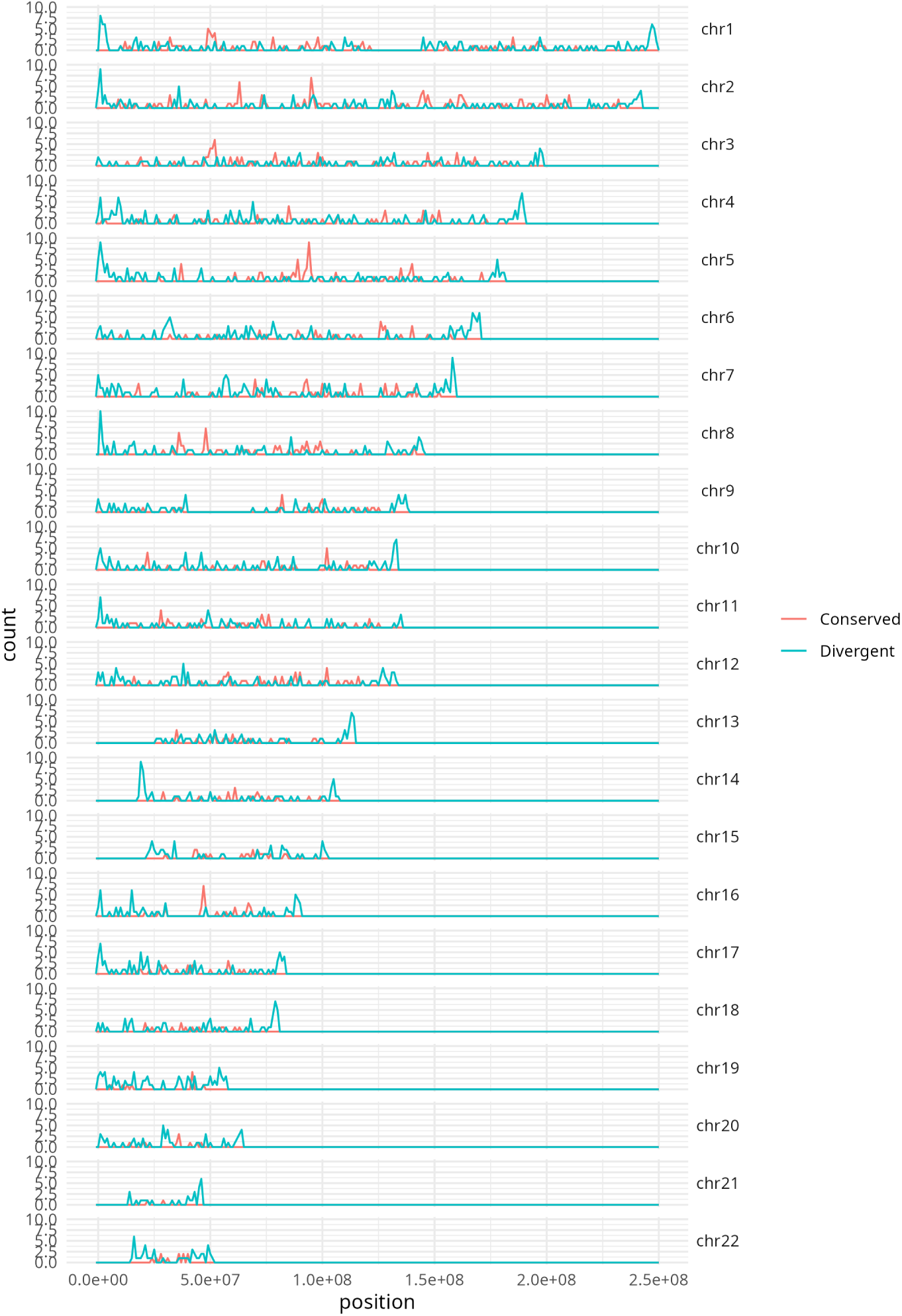
Density of conserved and divergent regions for bin size of 100k.

### C.5 Gene set enrichment

The gene set enrichment results for conserved regions are given in Figure 16. The first figure shows significantly enriched gene sets in the molecular function (MF), biological process (BP), and cellular compartment (CC) ontologies. The y-axis give the p-value.

**Figure 16:**
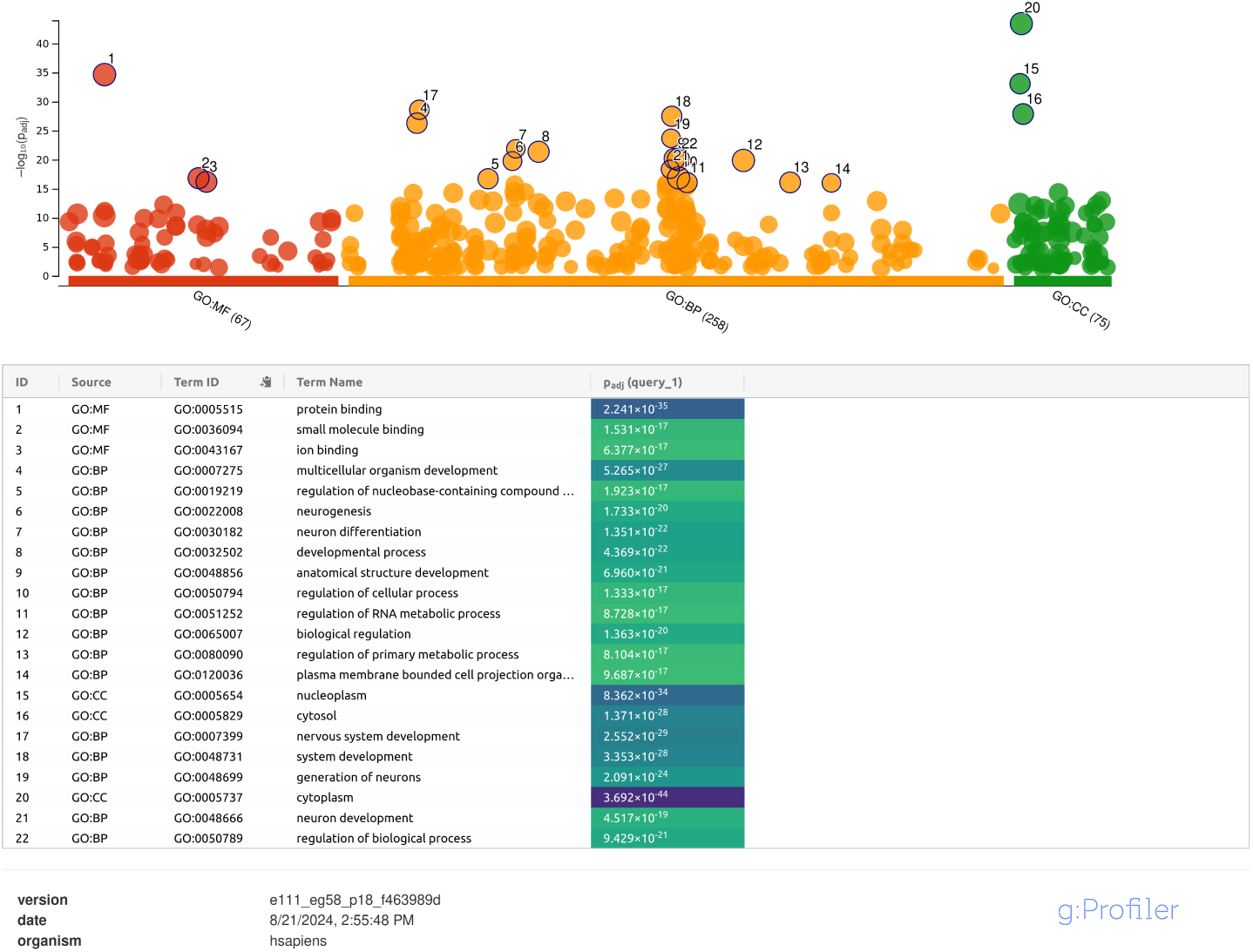
Gene Ontology results for conserved regions, computed with a window size of 10k.

Enriched gene sets are very vague, and shared in most species.

The gene set enrichment results for divergent regions are given in Figure 17. Here, some genes related to immunity can be found in all three ontologies.

**Figure 17:**
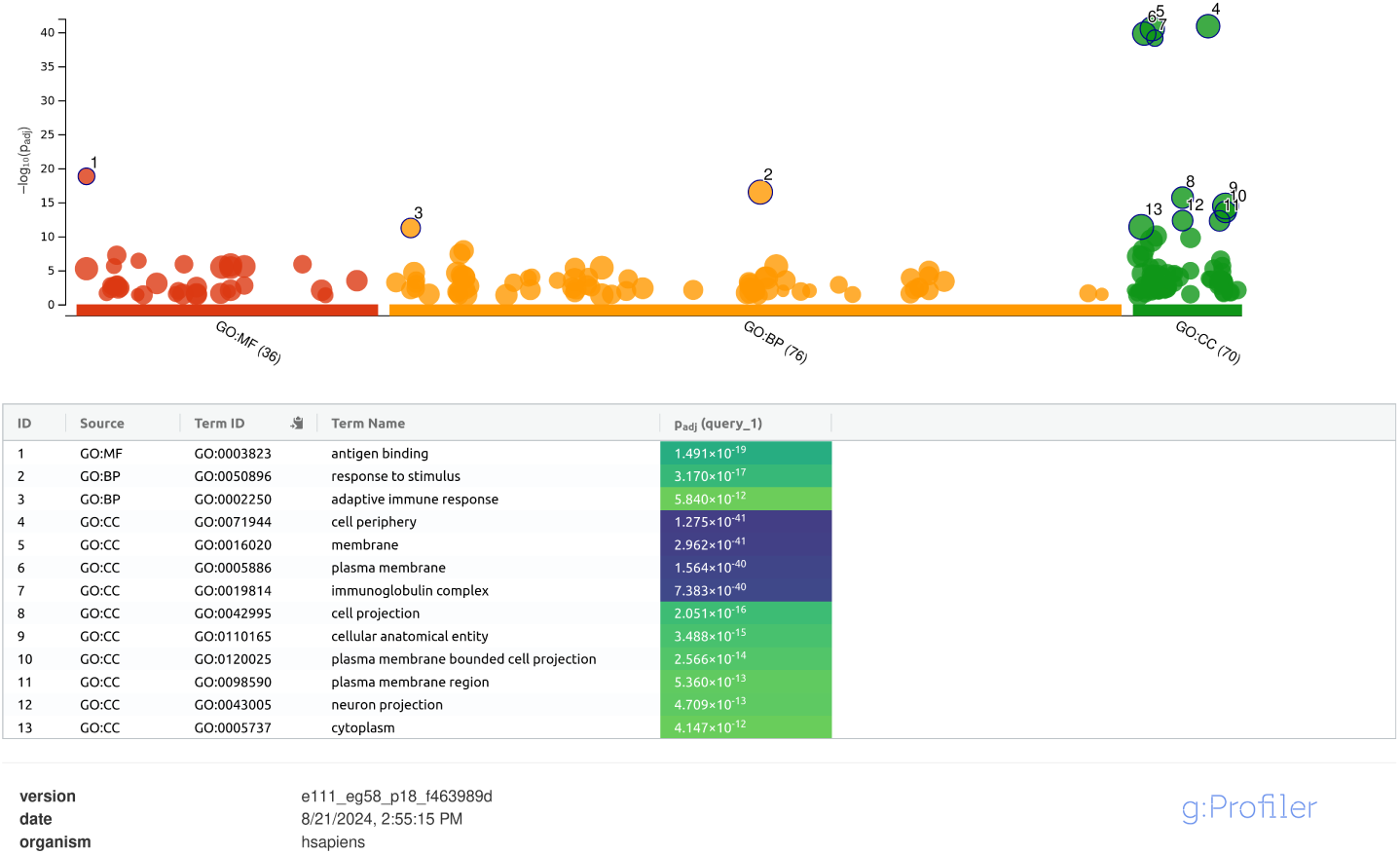
Gene Ontology results for divergent regions, computed with a window size of 10k.

